# Modelling alcohol consumption in rodents using two-bottle choice home cage drinking and optional lickometry-based microstructural analysis

**DOI:** 10.1101/2024.07.19.604367

**Authors:** Shikun Hou, Nathaly M Arce Soto, Elizabeth J Glover

## Abstract

Two-bottle choice home cage drinking is one of the most widely used paradigms to study ethanol consumption in rodents. In its simplest form, animals are provided with access to two drinking bottles, one of which contains regular tap water and the other ethanol, for 24 hr/day with daily intake measured via change in bottle weight over the 24 hr period. Consequently, this approach requires no specialized laboratory equipment. While such ease of implementation is likely the greatest contributor to its widespread adoption by preclinical alcohol researchers, the resolution of drinking data acquired using this approach is limited by the number of times the researcher measures bottle weight (e.g., once daily). However, the desire to examine drinking patterns in the context of overall intake, pharmacological interventions, and neuronal manipulations has prompted the development of home cage lickometer systems that can acquire data at the level of individual licks. Although a number of these systems have been developed recently, the open-source system, LIQ HD, has garnered significant attention in the field for its affordability and user friendliness. Although exciting, this system was designed for use in mice. Here, we review appropriate procedures for standard and lickometer-equipped two-bottle choice home cage drinking. We also introduce methods for adapting the LIQ HD system to rats including hardware modifications to accommodate larger cage size and a redesigned 3D printed bottle holder compatible with standard off-the-shelf drinking bottles. Using this approach, researchers can examine daily drinking patterns in addition to levels of intake in many rats in parallel thereby increasing the resolution of acquired data with minimal investment in additional resources. These methods provide researchers with the flexibility to use either standard bottles or a lickometer-equipped apparatus to interrogate the neurobiological mechanisms underlying alcohol drinking depending on their precise experimental needs.

**SUMMARY:** This protocol describes a standard intermittent-access two-bottle choice home cage drinking paradigm to model alcohol consumption in rats. In addition, it provides step-by-step instructions to augment the standard protocol with a DIY lickometer system that enables microstructural analysis of drinking behavior.

## INTRODUCTION

Although moderate alcohol use has historically been associated with moderate health benefits, more recent large-scale comprehensive studies have revealed that no amount of alcohol consumption is safe^1, 2^. In fact, alcohol use is the seventh leading risk factor for death and disability globally^1^ with individuals who drink even small amounts of alcohol having increased risk for cancers, infectious disease, and injury^1, 3^. In the United States, deaths from alcohol use increased by almost 30% between 2016 and 2021^4^. Importantly, risk for death and disability increases monotonically with increased consumption^1^. Likewise, patterns of alcohol consumption are well known to impact both risk for and severity of alcohol use disorder (AUD)^5–7^.

Although studies in humans have provided valuable insight into the neurobiology of alcohol use and misuse, the experimental control afforded by animal models is crucial for an in-depth understanding of the mechanisms underlying drinking behavior and risk for heavy drinking. These models are also valuable tools for the development of treatments aimed at reducing uncontrolled consumption. Two-bottle choice home cage drinking is one of the most widely used preclinical paradigms to study alcohol consumption in rodents. This is due, in large part, to its ease of implementation as it allows for the measurement of total voluntary alcohol drinking and preference without the need for specialized research equipment or complex analyses. Using this approach, researchers can collect measurements at their desired interval (e.g., hours, days, weeks) by calculating change in bottle weight from the beginning to the end of a drinking session. Variations on this basic method have been used by many in the field to facilitate low to moderate to binge levels of alcohol intake over both short and long periods of time. For example, continuous and intermittent two-bottle choice procedures have been used to facilitate low and moderate levels of alcohol intake, respectively, in both mice^8^ and rats^9–11^. The same paradigms can promote high levels of intake in alcohol preferring strains^10, 12^. Alternatively, an adaptation of this method that limits alcohol access to 2-4 hours daily beginning approximately three hours after the start of the dark cycle has been shown to engender binge drinking in mice (i.e., Drinking in the Dark)^13,14^. Additional variations on these methods have been used in conjunction with methods that facilitate alcohol dependence (i.e., chronic intermittent ethanol vapor exposure) to examine escalation of voluntary alcohol drinking during withdrawal^15–20^.

Although widely adopted by the field, the standard approach of measuring alcohol intake by change in bottle weight is limited in resolution. Increasing desire to examine drinking patterns in addition to levels of overall intake has prompted the development of a number of lickometer-equipped systems of varying complexity that can capture data at the level of individual licks. Among the first was the development of a two-bottle system equipped with photobeam lick detection designed for use in mice^21^. This system was subsequently adapted by a separate group for use in rats^22^. In the years since, variations on this theme have been used to develop similar systems contained within custom rodent housing^23^, that allow for lick detection of individual socially housed animals^24, 25^, and that capture both feeding and drinking behavior^26^. Among these, LIQ HD^27^ has garnered significant attention for offering a system for use in mice that parallels the ease of implementation and affordability of the system developed by Godnyuck et al.^21^ but uses a capacitance sensor, which the authors demonstrate affords greater precision over photobeam lick detection. The authors further demonstrate the ability to use this system in conjunction with a continuous access two-bottle choice home cage drinking paradigm to capture measures of alcohol drinking microstructure in addition to measures of overall daily intake^27^.

Here, we describe procedures for two-bottle choice home cage alcohol drinking using both standard and lickometer-equipped systems. We also introduce methods for building and implementing LIQ HDR – an adaptation of LIQ HD^27^ for use in rats that includes hardware modifications to accommodate larger cage size and a redesigned 3D printed bottle holder compatible with standard off-the-shelf bottles. These methods provide researchers with the flexibility to use either a standard or lickometer-equipped approach depending on their experimental resources and data collection needs.

**Figure 1:**
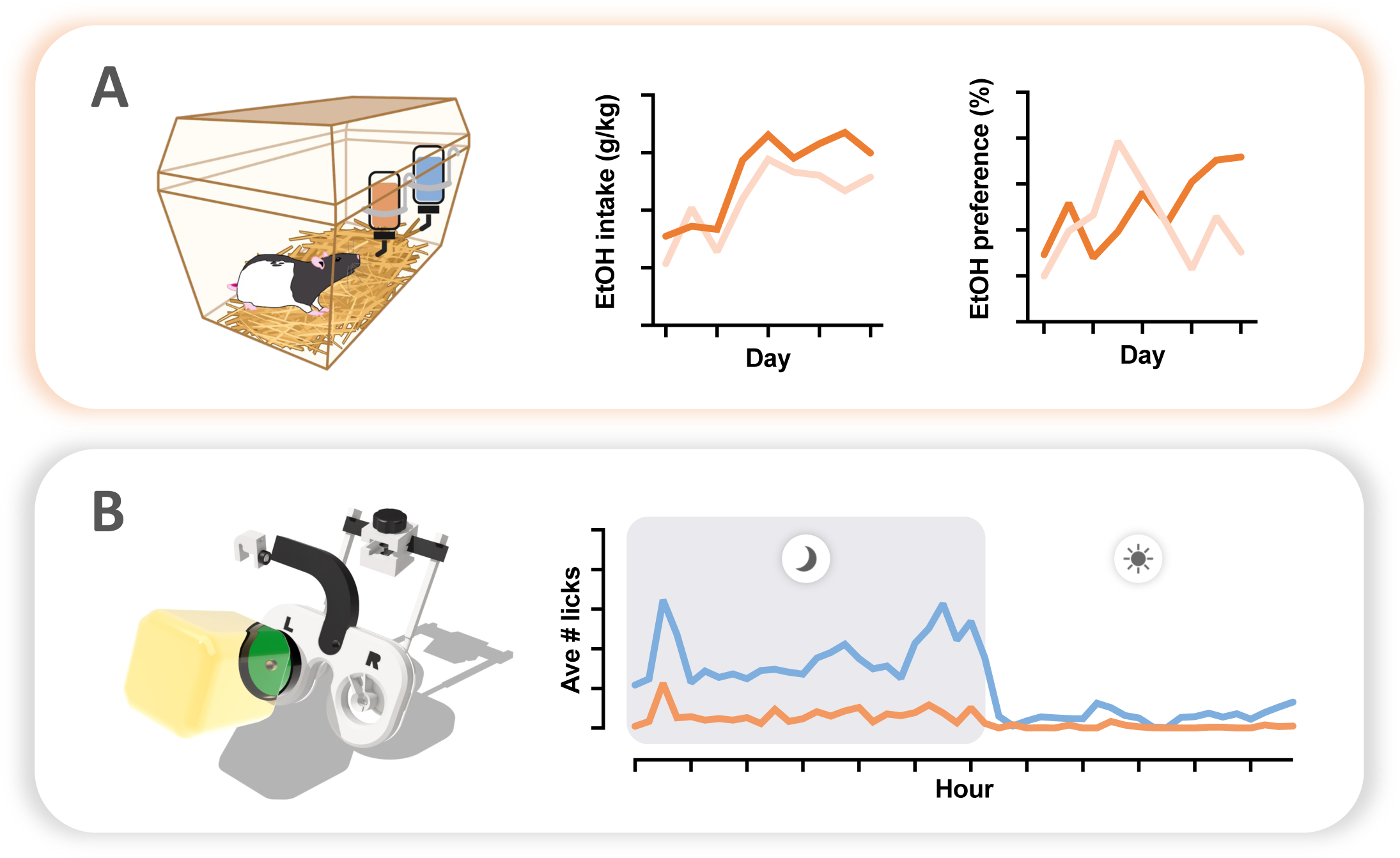
Two-bottle choice home cage drinking. **(A)** Researchers can measure voluntary home cage ethanol intake and preference (relative to water) using a standard approach that calculates intake based on change in bottle weight. **(B)** By using LIQ HDR, a low-cost DIY home cage lickometer system designed for rats, researchers can acquire the same measures as in the standard two-bottle choice procedure while also capturing high-resolution drinking microstructure during the entire recording period.

## PROTOCOL

NOTE: Below is a description of materials needed and step-by-step instructions to capture home cage drinking using the intermittent-access two-bottle choice drinking procedure adapted from Simms et al.^10^. In this procedure, rats are provided 24 hr access to two drinking bottles seven days a week. On Mondays, Wednesdays, and Fridays (MWF) rats receive one bottle containing water (H_2_O) and another bottle containing ethanol (EtOH). These bottles are removed on Tuesdays, Thursdays, and Saturdays (TRS) and replaced with two bottles of water. Rats drink from the same two bottles of water on Saturdays and Sundays. Note that these methods can easily be adjusted for alternate schedules of ethanol access (e.g., continuous, binge, etc) if desired. Separate instructions are provided for standard and lickometer-equipped approaches. All representative data was collected using adult Long Evans rats with the approval of the University of Illinois Chicago Institutional Animal Care and Use Committee and in accordance with the NIH Guidelines for the Care and Use of Laboratory Animals.

### 1. Preparation

NOTE: Prepare the following before beginning your experiment.

1. Using a sharpie, label drinking bottles as H_2_O or EtOH. Then label with unique identifier for each rat. NOTE: Each rat should be assigned one H_2_O and one EtOH bottle for MWF and two H_2_O bottles for TRS. Optionally, use different color tape to identify H_2_O and EtOH bottles to help avoid potential error during bottle change
2. Fill H_2_O bottles with drinking water one day prior to the experiment start date.
3. Prepare EtOH solution by combining 190 proof EtOH with drinking water (20% v/v) one day prior to the experiment start date. CAUTION: It is critical that the EtOH used is non-denatured as denatured EtOH is poisonous.
4. Prepare an empty cage on the same shelving as the rats participating in the experiment. This cage will be used to calculate spillage.
5. Equip cages with LIQ HDR lickometer system if using (see **Section 4**, **Figures 2-6, & Supplemental Figure 1).**
6. Determine the precise time of day for bottle change prior to the onset of the study. Bottles should be replaced at the same time every day in order to obtain an accurate measurement of 24 hr intake. NOTE: Previous research has shown that rodents exhibit the greatest degree of consumption early after the onset of the dark cycle^13, 28^. Therefore, in order to take advantage of this highly active period, it is recommended that bottle change occur 30-60 min after lights are turned off.

### 2. Daily procedures

1. On the first day of the experiment, do the following:

1. Remove standard water bottles from cages.
2. *Optional:* Rats often jostle the cage top as they are foraging for food, which could cause spillage that cannot be corrected for by the empty spillage cage. The use of food cups inside the cage can alleviate this concern. If using, remove food from the cage top now, add a food cup inside the cage, and fill with chow.
3. Indicate to care and husbandry staff as applicable that research personnel will provide food and water for the duration of the experiment.
4. *Optional:* Cage changes can be disruptive to data collection. We recommend that research personnel perform cage changes in tandem with obtaining body weight measurements. Researchers choosing to do this should inform their care and husbandry staff accordingly.
2. On subsequent days, follow the procedures outlined below: NOTE: Remember that any movement of cages and bottles can create spillage. Therefore, it is important to be gentle with movements and use the same methods of applying and removing bottles on the spillage cage as are used for rat cages.

1. Enter the room prior to bottle change time ensuring that you have enough time to carry out the steps below and place new bottles on cages at the appropriate session start time.
2. Remove filter tops from cages. If MWF, proceed to **step 2.3**. If TRS, proceed to **step 2.9**. On Mondays, Wednesdays, and Fridays, perform the following procedures:
3. Remove TRS bottles from cage tops. NOTE: Typically, water consumption is not recorded or reported for TRS, but researchers can optionally collect these data by obtaining on and off bottle weights as described below in **step 2.4** & **2.10** if desired.
4. Weigh MWF bottles (one EtOH and one H_2_O). Record as bottle start weight for today’s drinking session.
5. Provide additional food to cage top or food cups if necessary.
6. At designated session start time, place bottles on each rat’s cage top with tape facing up so that rats do not chew the bottle label. NOTE: The side for the EtOH bottle should be alternated after each drinking session in order to avoid the development of potential side preference, which can inadvertently skew consumption data.
7. *Optional:* Previous work has shown that many rats exhibit binge-like drinking at the onset of the drinking session^11, 29, 30^. Most researchers capturing binge drinking during this time report intake during the first 30-60 min of the drinking session. To obtain intake data during this potential binge episode:
  1. Set a timer after bottles are applied to the last cage
  2. Leave the room undisturbed during the timed period
  3. Re-enter the room when timer goes off
  4. Remove EtOH and H_2_O bottles and record bottle weights
  5. Return EtOH and H_2_O bottles to cages
  NOTE: This step is not required to capture binge episodes if using the lickometer-equipped system.
8. Return filter tops to cages and leave the animal room. On Tuesdays, Thursdays, and Saturdays, perform the following:
9. Remove MWF bottles from cages.
10. Weigh each bottle. Record as bottle end weight for the MWF drinking session. NOTE: Optionally cover EtOH bottle sippers to prevent evaporation.
11. Obtain and record a body weight for each rat. NOTE: If cage changes are being performed by research personnel, designate one day weekly during TRS duties to perform this task. Place each rat into a clean cage after obtaining body weight.
12. Provide additional food to cage top or food cups if necessary.
13. At designated session start time, place TRS H_2_O bottles on each rat’s cage top with tape facing up so that rats do not chew the bottle label.
14. Return filter tops to cages and leave the animal room.

### 3. Maintenance

1. Wash bottles and refresh fluids on a regular weekly or biweekly basis in accordance with your institution’s animal care and husbandry procedures NOTE: Care should be taken to perform these duties in a manner that does not interrupt drinking sessions. For example, TRS H_2_O bottles can be washed and refilled after removal on MWF. Similarly, MWF H_2_O and EtOH bottles can be washed and refilled after removal on TRS.

### 4. Adding lickometers to two-bottle choice experiments

NOTE: Researchers interested in capturing high resolution drinking data can modify the above procedure by adding lickometers to the bottles. LIQ HD^27^ is a low-cost DIY system that allows for this type of data collection in mice. The procedures outlined below describe the modifications necessary to adapt LIQ HD for use in rats (referred to as LIQ HDR). See **Table of Materials** for a list of all commercially sold and custom 3D-printed parts necessary to construct one lickometer system, which can capture two-bottle choice data for up to 18 rats. The 3D print .stl files and instructions can be found at the author’s website (www.ejgloverlab.com/3dprints).

1. Setting up capacitive sensor units NOTE: Each LIQ HDR system consists of three capacitive sensor units, which together capture lick data from 36 bottles in parallel corresponding to 18 single-housed rats. Each sensor unit consists of one QwiicBus communication board and one 12-key capacitive sensor board, which will connect to lickometers from six separate cages via six pairs of 2-pin cables that are color-coded for each lickometer (**Figure 2A**).

**Figure 2:**
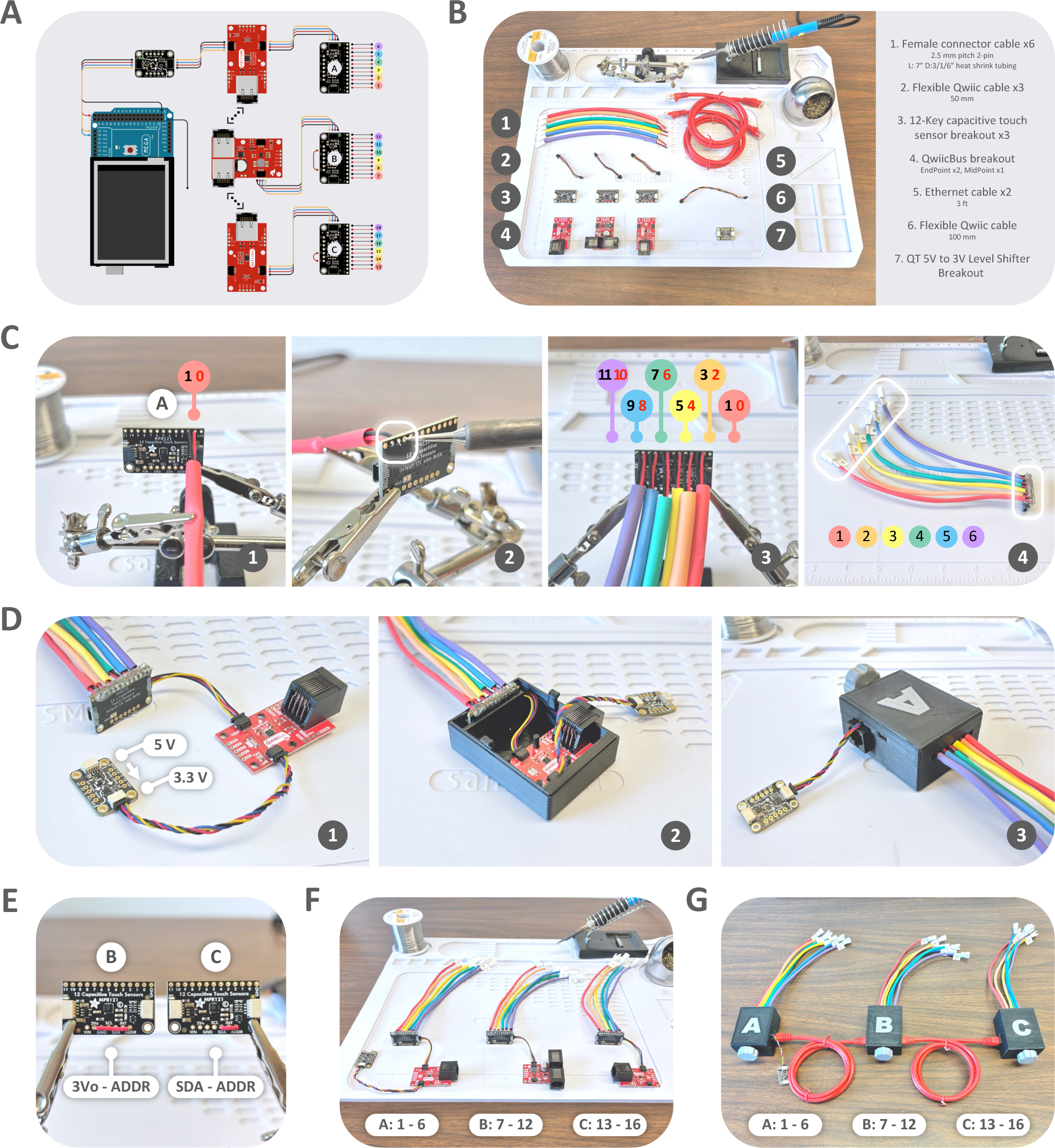
Constructing capacitive sensor units. **(A)** Electronic parts and wiring diagram. **(B)** Materials for constructing three capacitive sensor units required build one LIQ HDR system. **(C)** Solder female connector cables for cages #1-6 to board A pin #0-11 and secure the solder joints with hot glue. Color code cables with colored heat shrink tubing and label each with a unique lickometer ID number. **(D)** Connect board A and the 5V-to-3V level shifter to a QwiicBus EndPoint with Qwiic cables (1), and house board A and QwiicBus EndPoint in the 3D printed casing (2-3). **(E)** Change the I^2^C addresses of boards B and C by soldering a jumper wire from the ADDR pin to the 3Vo and SDA pins, respectively. **(F)** Solder female connector cables to sensor boards B and C, label with the corresponding lickometer IDs, and connect board B to a QwiicBus MidPoint and board C to an EndPoint with Qwiic cables. **(G)** House sensor units B and C in their respective 3D print cases and daisy-chain all three units with Ethernet cables.
  1. Gather the materials required for building all three capacitive sensor units as shown in **Figure 2B**, along with required soldering tools.
  2. To start building sensor unit A, cover a 2-pin female connector cable with a 7” long, red heat-shrink tubing, insert its red and black wire ends to pin #0 and #1 on the sensor board (**Figure 2C-1**), and solder in place (**Figure 2C-2**). Make sure that the cable is at the front of the sensor board as shown in order to arrange the cages in numerical order from left to right in the vivarium without overlapping the cables and potentially disrupting recording (see **step 6** & **Figure 6**)
  3. Cover five more cables with distinctly colored heat shrink tubing and solder to pin #2-11 as shown (**Figure 2C-3**). Make sure all red wires are soldered to even numbered pins, which will connect to all bottles on the left side, and all black wires are soldered to odd numbered pins, which will connect to all bottles on the right side.
  4. Cover the solder joints with hot glue on all sides for protection (**Figure 2C-4**).
  5. Label the connectors in numerical order providing a unique ID for each lickometer (**Figure 2C-4**).
  6. To assemble sensor unit A, connect sensor board A to a QwiicBus EndPoint with a 50 mm Qwiic cable, and connect the EndPoint to the 3.3 V output of the 5V-to-3V lever shifter with a 100 mm Qwiic cable (**Figure 2D-1**). Although both the sensor board and QwiicBus communication board have two identical Qwiic connectors, it is still important to make sure that the wiring is exactly as shown in (**Figure 2D-1** & **Figure 2F**) in order to use the 3D printed sensor cases and our suggested vivarium setup (see **step 6** & **Figure 6**). House sensor unit A inside the 3D printed case (**Figure 2D-2**). Gently close the top taking care not to crush the level shifter cable (**Figure 2D-3**).
  7. Modify the I^2^C addresses of sensor board B and C by soldering a jumper wire from the ADDR pin to the 3Vo or SDA pin as indicated (**Figure 2E**).
  8. Solder six color-coded female connector cables to sensor board B and C using the same order used for board A. Label each connector with the corresponding lickometer ID number. Connect board B to the QwiicBus MidPoint and C to a MidPoint with 50 mm Qwiic cables as shown (**Figure 2F**).
  9. House sensor unit B and C in their respective 3D printed cases. Daisy-chain all three sensor units with Ethernet cables to complete sensor setup (**Figure 2G**).
2. Set up the Arduino interface NOTE: The LIQ HDR Arduino interface (**Supplemental Figure 1A**) is the same as described for LIQ HD^27^ with very minor modifications. The procedures outlined below provide a general overview of the steps necessary to set up the interface with modifications described in steps 10 and 12. Researchers are referred to the original protocol and the video tutorial provided on the author’s GitHub page for more detailed step-by-step instructions^27^.

**Supplemental Figure 1:**
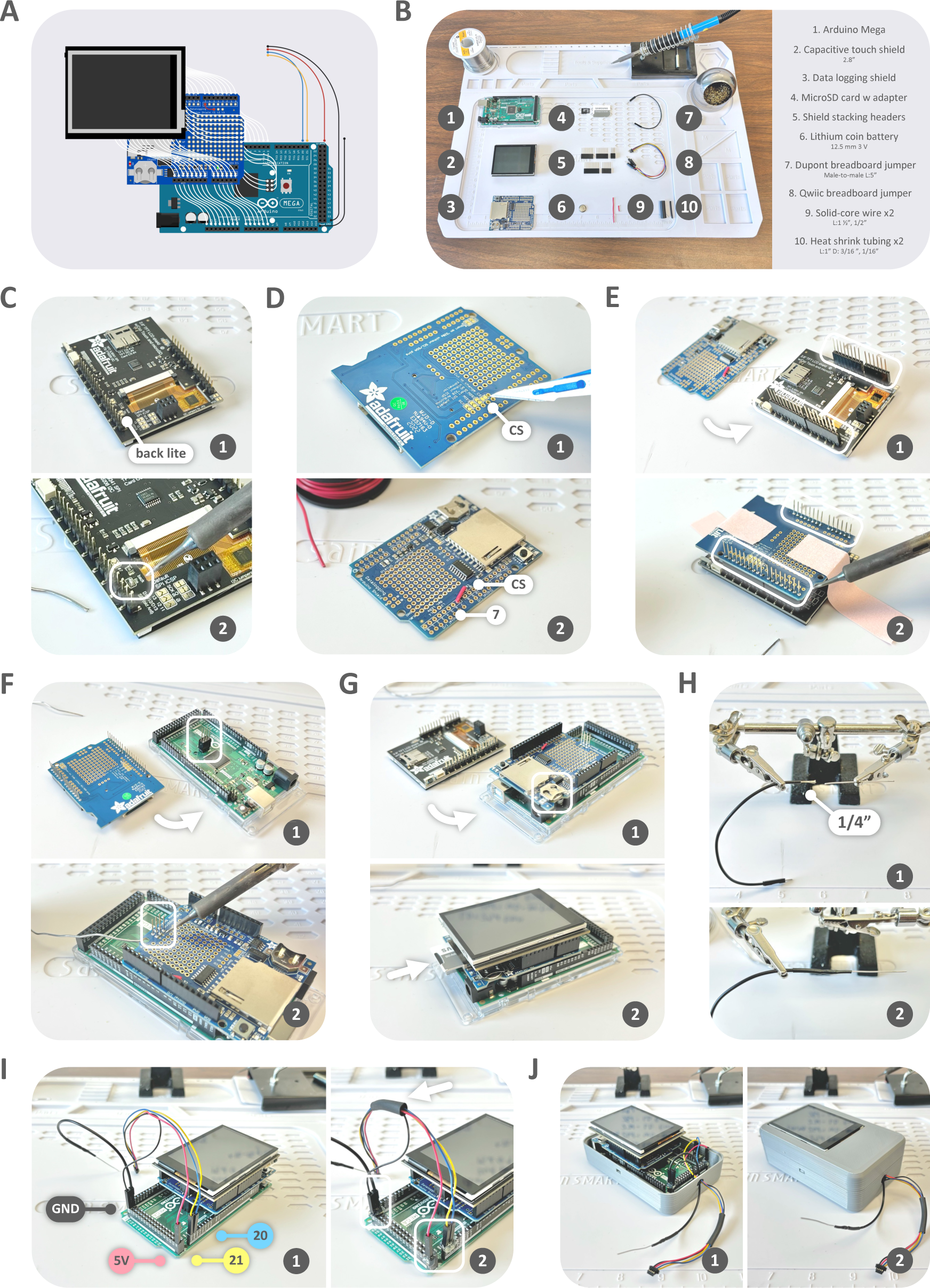
Set up the Arduino interface. **(A)** Electronic parts and wiring diagram**. (B)** Materials for building the Arduino interface required for one LIQ HDR system. **(C)** On the back side of the touchscreen shield, add a solder jumper across the back lite solder pads. **(D)** On the back side of the data logging shield, sever the jumper pad trace of the CS pin (1). Solder a jumper wire from the CS pin to pin #7 (2). **(E)** Solder header pins to the data logger using the touchscreen shield as the base. **(F)** Stack the remaining 2x4 header to the Arduino Mega (1), stack the data logger on the Arduino Mega and solder the header in place (2). **(G)** Insert the coin battery into the data logger (1), stack the touchscreen onto the data logger, and insert the SD card into the data logger (2). **(H)** To make the extra ground wire, cut off one connector from the breadboard jumper and solder the end to a stripped solid-core wire. **(I)** Connect the Qwiic jumper wires and extra ground wire to the Arduino MEGA as shown (1), and secure with hot glue (2). **(J)** House the Arduino interface in the custom 3D printed casing.
  1. Gather the materials and soldering tools required to build the Arduino interface (**Supplemental Figure 1B**).
  2. On the back side of the touchscreen shield (**Supplemental Figure 1C-1**), create a solder jumper across the back lite solder pads (**Supplemental Figure 1C-2**).
  3. On the back side of the data logging shield, sever the jumper pad trace of the CS pin (**Supplemental Figure 1D-1**).
  4. On the front side of the data logger, solder a short jumper wire from the CS pin to pin #7 (**Supplemental Figure 1D-2**).
  5. To install the shield stacking headers, first stack them to the back of the touchscreen shield (**Supplemental Figure 1E-1**). Stack the data logger to the touchscreen, use tape to stabilize, and solder all the header pins to the data logger (**Supplemental Figure 1E-2**). Then remove the touchscreen.
  6. Stack the remaining 2x3 header to the Arduino Mega (**Supplemental Figure 1F-1**), then stack the data logger onto the Arduino before soldering the header pins in place (**Supplemental Figure 1F-2**).
  7. Insert the coin battery into the data logger (**Supplemental Figure 1G-1**).
  8. Stack the touchscreen onto the data logger (**Supplemental Figure 1G-1**).
  9. Insert the microSD card along with its adapter into the data logger (**Supplemental Figure 1G-2**).
  10. To make the extra grounding wire, cut off one connector from the breadboard jumper wire and strip the end to ∼1/4” (**Supplemental Figure 1H-1**). Solder a strand of stripped solid-core wire to the jumper wire and cover the solder joint with heat shrink tubing (**Supplemental Figure 1H-2**).
  11. Plug the Qwiic male jumper and extra grounding wire to the Arduino Mega as shown (Black – GND, Red – 5V, Yellow – pin #21, Blue – pin #20, **Supplemental Figure 1I-1**). Secure the connection with hot glue (**Supplemental Figure 1I-2**). Optionally, wrap the Qwiic jumper wires with heat shrink tubing (**Supplemental Figure 1I-2**).
  12. House the fully assembled Arduino interface in the 3D printed case with the Qwiic and grounding wire threaded through the opening at the top (**Supplemental Figure 1J-1&2**).
  13. Follow the original protocol for Arduino interface software installation^27^.
3. Build LIQ HDR lickometers NOTE: The custom 3D print that holds the bottles is designed for use with a standard rodent drinking bottle with a sipper tube diameter of 8 mm (see **Table of Materials**, **Figure 3A**). This bottle holder can easily be adapted for use with similar bottles and sipper tubes with minor or no modifications.

**Figure 3:**
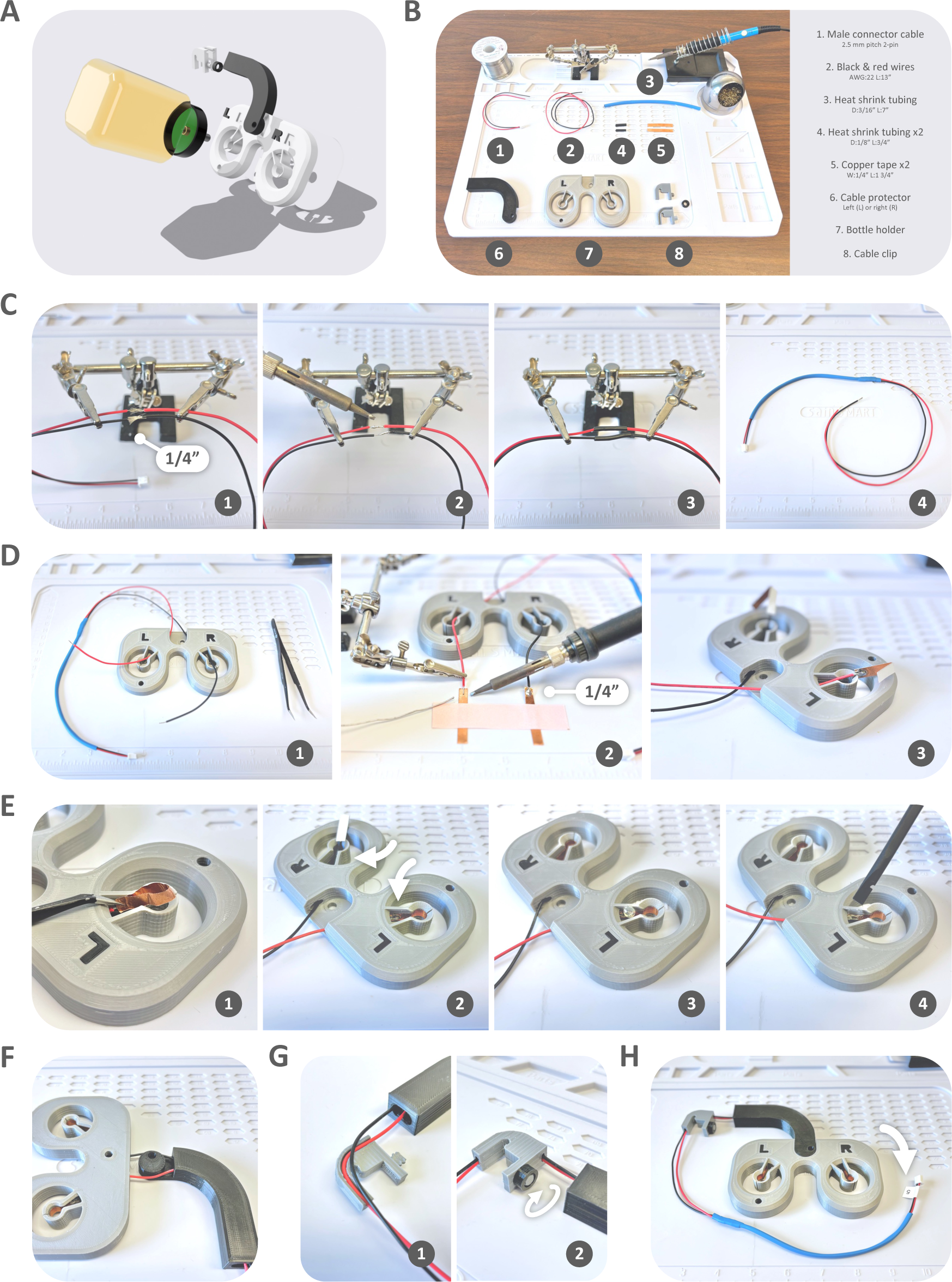
Build the LIQ HDR lickometer. **(A)** 3D rendering of the lickometer in-cage components. **(B)** Materials for setting up one two-bottle lickometer. Note that one LIQ HDR system accommodates 18 two-bottle lickometers. **(C)** Extend the length of the male connector cable with a pair of black and red wires and color code with heat shrink tubing. **(D)** Thread the black and red wires through the left (L) and right (R) openings on the bottle holder (1) and solder the ends to the copper tapes (2-3). **(E)** Adhere the copper tapes to the inner walls of the sipper brackets (1-2) and reinforce with hot glue (3-4). **(F)** Thread the cable through the cable protector. **(G)** Install the cable clip. **(H)** Label the connector with matching lickometer ID number.
  1. Gather the materials, 3D prints, and soldering tools required for building one lickometer as shown in **Figure 3B**. Note that each LIQ HDR system accommodates a maximum of 18 lickometers for 36 bottles.
  2. To extend the male connector cable, strip both of its ends to ∼1/4” in length, as well as the extension black and red wires (**Figure 3C-1**).
  3. Solder the extension wires to the connector cable (**Figure 3C-2**) and cover the solder joints with heat shrink tubing (**Figure 3C-3**). Cover the male connector cable with heat shrink tubing that is colored matched to a corresponding female connector cable on the sensor unit (**Figure 3C-4**).
  4. To be consistent with the sensor board wiring (**Figure 2**), thread the red and black wire through the left and right internal cable passage of the bottle holder, respectively (**Figure 3D-1**).
  5. Solder each wire to the very end of the 1 ¾” conducting copper tape (**Figure 3D-2**) and fold the copper tapes in half as shown (**Figure 3D-3**).
  6. Unpeel the copper tape and use a pair of forceps to carefully maneuver it inside the sipper bracket (**Figure 3E-1**) before firmly adhering it to the inner walls of the sipper bracket (**Figure 3E-2**). Make sure that the solder side of the copper tape is adhered to the wall closer to the midline of the bottle holder as indicated (**Figure 3E-2**). Push a sipper tube through the bracket to press the tape firmly against the wall and prime the bracket.
  7. To reinforce the solder side of the copper tape, which is not as adhesive and tends to peel off, apply a dollop of hot glue to the solder joint (**Figure 3E-3**). Smear the glue to cover both the copper tape and any adjacent exposed portion of the 3D print (**Figure 3E-4**).
  8. Thread the cable through the internal passage of the cable protector (**Figure 3F**). NOTE: Be sure to select the appropriate cable protector (left- or right-facing) for each lickometer in order to use the suggested vivarium setup (see **step 6** & **Figure 6**). Lickometers 5-6, 11-12, and 17-18, which are depicted in blue- or purple-colored cables in the current set up, require a left-facing cable protector (**Figure 4D-1**), whereas all other lickometers require a right-facing cable protector (**Figure 4D-2**).

**Figure 4:**
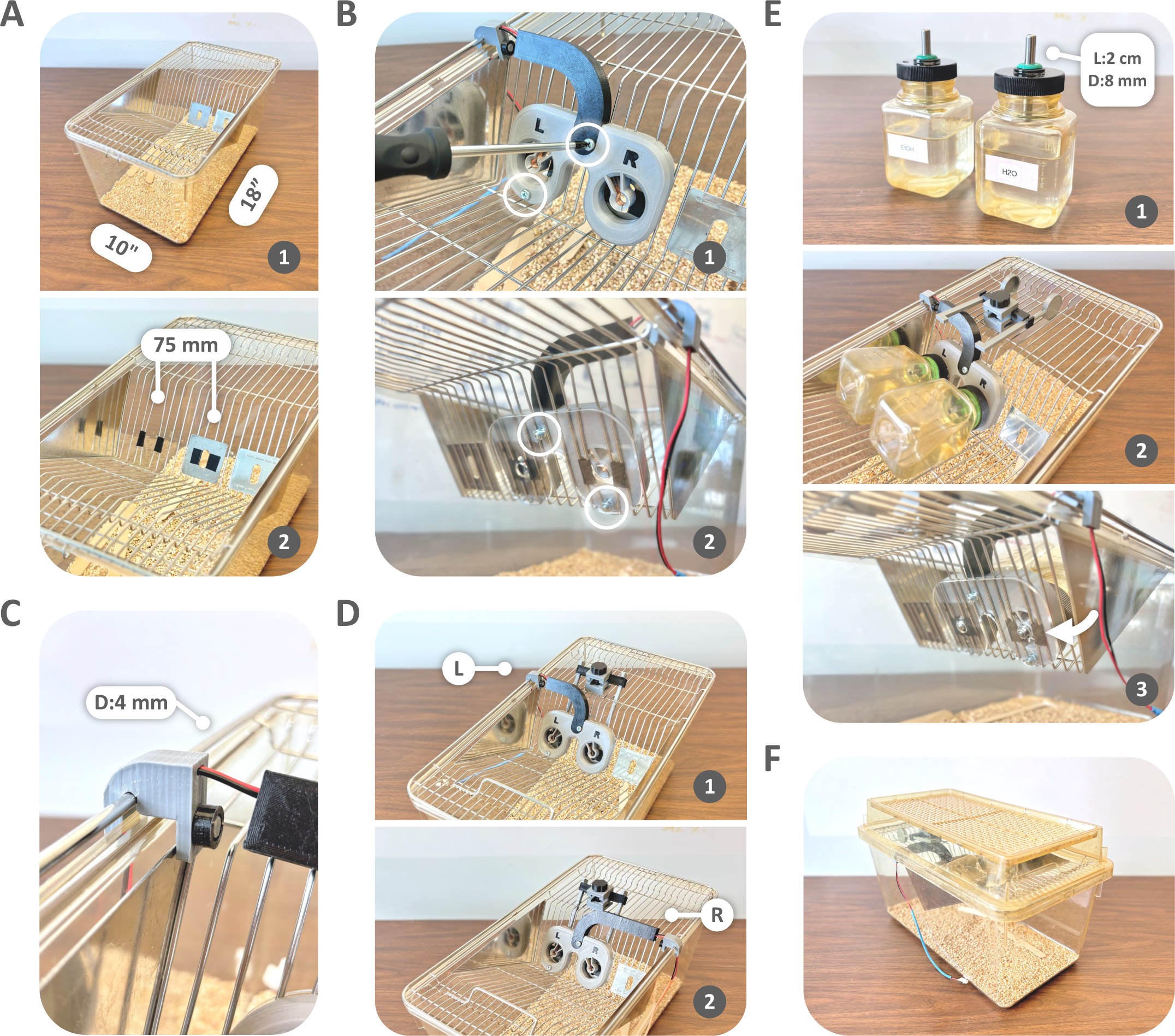
Lickometer installation. **(A)** Use electrical tape to insulate the metal rods around the sipper openings on the metal cage top. **(B)** Install the lickometer and acrylic panel to the cage top with two sets of M5 screws and nuts. **(C)** Secure the cable clip to outer-most rod of the wire cage top. **(D)** Make sure to use the appropriate cable protector (left L or right R) for each cage. **(E)** Make sure only ∼2cm of the bottle sippers are exposed (1), so that when the bottles are placed on the cage (2) only the very tips of the sippers extend beyond the acrylic panel and are accessible to rats. **(F)** The filter top should fit over the cable clip without disturbing the cable itself.
  9. To install the cable clip, fit one wire in the groove of one half of the cable clip first (**Figure 3G-1**), then fit the other half over the second wire and tightly close both halves with the nut (**Figure 3G-2**).
  10. Label the male connector with the corresponding lickometer ID number (**Figure 3H**).
  11. Repeat these steps until all 18 lickometers are built.
4. Install LIQ HDR lickometers NOTE: LIQ HDR is designed for installation on a standard shoebox rat cage with a metal wire top (**Figure 4A-1**).
  1. Identify a pair of sipper openings that are ∼75 mm apart in center distance on the wire cage top in order to accommodate the bottle holder. Use a pair of pliers to open slightly the metal rods around the sipper openings. Insulate these rods with electrical tape in order to prevent metal-to-metal contact between the sippers and the cage top (**Figure 4A-2**).
  2. Install the lickometer and the 1/8” thick laser acrylic panel to the cage top with two sets of M5 screw and nut (**Figure 4B**). Make sure that the sipper openings on the lickometer and acrylic panel align, and the sippers are not in contact with any part of the cage top.
  3. Secure the cable clip to the outer-most rod (∼4 mm in diameter) of the wire cage top (**Figure 4C**). Adjust the cable clip if needed to make sure that the cage top rests snuggly on the cage bottom and that the cable is not being pulled or bent under too much tension.
  4. Make sure that the correct cable protector is being used for each lickometer (**Figure 4D** & see **step 3.8**).
  5. Adjust the sipper tubes as necessary so that only ∼2 cm of the sipper extends from the cap (**Figure 4E-1**). To install bottles, insert the sipper tube through the sipper bracket of the bottle holder until it stops (**Figure 4E-2**). Only the very tip of the sipper tube should extend beyond the acrylic panel (**Figure 4E-3**). It is crucial that no other part of the sipper tube is exposed in order to avoid false positive licks from non-consummatory behaviors, such as the rat touching or playing with the sipper tube. NOTE: A thicker acrylic panel or a stack of multiple 1/8” panels can be used to fully enclose sipper tubes of longer lengths for experiments that require that sipper tubes extend further into the cage, such as when measuring drinking in young rats that may have trouble reaching a short sipper.
  6. A filter top should be able to rest snugly over the wire cage top and cable clip without disturbing the cable itself (**Figure 4F**).
5. Assemble & install sipper blockers NOTE: The sipper blocker (**Figure 5A**) is designed to restrict access by the rat to the sipper during bottle changes and to ensure accurate lickometer calibration at the onset of each drinking session.

**Figure 5:**
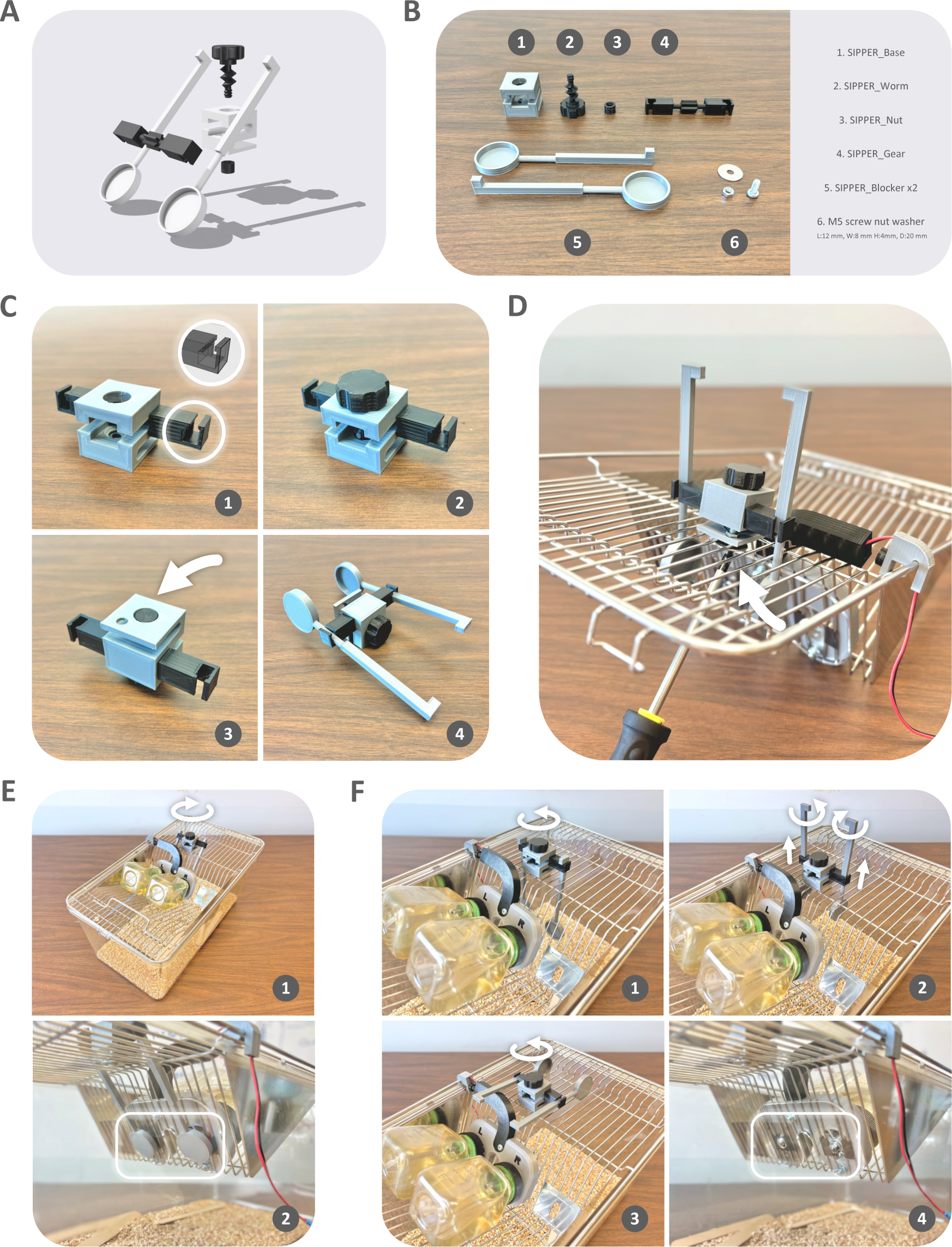
Sipper blocker installation and use. **(A)** 3D rendering of the sipper blocker. **(B)** Custom 3D printed parts for one sipper blocker. **(C)** Assemble the 3D printed parts. **(D)** Install the assembled sipper blocker onto the cage top with an M5 screw + nut + washer. **(E)** Turn the dial clockwise to block access to the sippers. **(F)** Turn the dial counterclockwise to remove blockers from the sippers (1), lift the blockers and rotate 90° as shown (2), and continue turning the dial counterclockwise until the blockers are above the cage top (3-4).
  1. Gather the 3D printed parts and an M5 screw, nut, and washer for one sipper blocker as shown in **Figure 5B**.
  2. Insert the SIPPER_Gear into the SIPPER_Base (**Figure 5C-1**). Make sure that the SIPPER_Gear is oriented exactly as shown in the 3D rendering inset.
  3. Insert the SIPPER_Worm into the opening at the top of the SIPPER_Base. Turn the worm clockwise while inserting until the head is flush with the base (**Figure 5C-2**).
  4. Screw the SIPPER_Nut to the end of the Worm from the bottom of the Base (**Figure 5C-3**). Use forceps to tighten the nut while turning the dial of the worm.
  5. Insert the round narrow necks of the SIPPER_Blockers into the slots of the Gear (**Figure 5C-4**).
  6. Install the assembled sipper blocker onto the cage top using the M5 screw set (**Figure 5D**). Make sure that both sipper blockers align with the sipper openings of the lickometer.
  7. To block access to the sippers, turn the worm dial clockwise to lower the blockers until they completely cover the sipper openings (**Figure 5E**).
  8. To provide access to the sippers, turn the dial counter-clockwise until the blockers are in a vertical position perpendicular to the cage top (**Figure 5F-1**). Manually pull up on each blocker arm until the heads are just below the wire cage top. Rotate the blockers 90° so that the heads are parallel to the metal rods on the cage top and the arms are locked in place (**Figure 5F-2**). Keep turning the dial counter-clockwise until both blockers are raised above the cage top (**Figure 5F-3**) and the sippers are fully accessible (**Figure 5F-4**).
6. Vivarium setup for high throughput data collection NOTE: The following vivarium setup is optimized to capture two-bottle choice drinking using a single Arduino interface connected to 18 standard shoebox rat cages. All electronic components, including the Arduino interface, sensor units, and various cables are secured to wall-mounted metal wire shelving that holds the cages (**Figure 6A**). Some modifications may be required to adapt to animal facilities and user needs.

**Figure 6:**
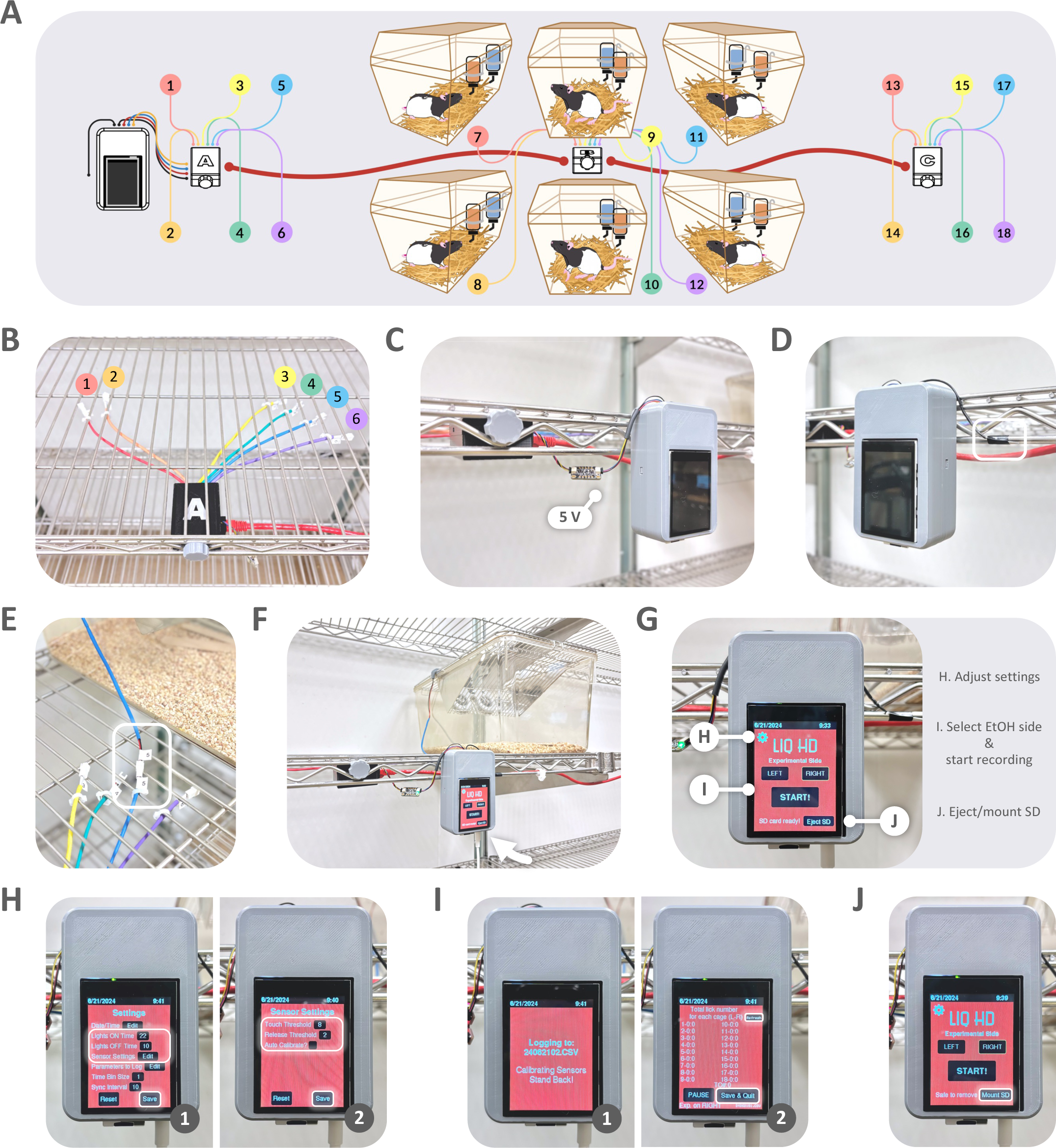
Vivarium setup and GUI operation. **(A)** Recommended vivarium setup and wiring diagram. **(B)** Mount each sensor unit to shelving and arrange and stabilize the sensor cables to the shelving. **(C)** Mount the Arduino interface to the shelving and connect its Qwiic jumper to the 5V-to-3V level shifter connected to sensor unit A. **(D)** Secure the extra ground wire with electric tape. **(E)** Connect the sensor cables to the corresponding lickometer cables. **(F)** Plug in the power supply to turn on the Arduino interface. **(G)** Main page of the GUI interface. **(H)** Tap Settings to adjust lights ON/OFF time to the vivarium light schedule (1) and tap Sensor Settings to adjust touch and release thresholds for rats (2). **(I)** To record data, select EtOH bottle side on the main page, tap START to calibrate the sensors (1) and start recording (2). Tap Refresh to update lick number shown on screen and tap Save & Quit to stop recording. **(J)** Tap Eject SD on the main page to remove the SD card and transfer data. Tap Mount SD after inserting SD card.
  1. Mount each sensor unit to the vivarium shelving using the custom 3D printed mounting screw (**Figure 6B**). Make sure that each sensor unit is mounted directly underneath the cage with the yellow-colored lickometer in order to avoid unnecessary cable tension (**Figure 6A**).
  2. Arrange and secure each female connector cable making sure that they do not overlap with each other or bend excessively at their solder ends, which could result in poor lick detection (**Figure 6B**). If metal rod shelving is used, this can be easily achieved by cable tying the connector cables to separate rods on the shelf as depicted.
  3. Mount the Arduino interface next to sensor unit A using the same 3D printed mounting screw and plug its Qwiic connector into the 5 V input port of the 5V-to-3V level shifter connected to sensor unit A (**Figure 6C**).
  4. Use electrical tape to secure the extra ground wire to a grounding source such as the metal shelving (**Figure 6D**).
  5. Arrange each cage based on the color and ID of the lickometer installed as per **Figure 6A** depiction in order to minimize cable overlap and tension.
  6. Securely connect all male lickometer connector cables to the matching female sensor connector cables (**Figure 6E**).
  7. Plug in the power supply to turn on the Arduino interface (**Figure 6F**).
  8. On the main GUI page (**Figure 6G**), tap the setting symbol to open the Settings page. Users can change “Lights ON and OFF time” based on to ensure that the interface display automatically dims when the dark cycle begins (**Figure 6H-1**).
  9. Tap “Edit Sensor Settings” (**Figure 6H-1**) to adjust “Touch Threshold” to 8 and “Release Threshold” to 2, which we have found to enable the most consistent lick detection for rats using the current setup (**Figure 6H-2**). Note, however, that users may need to experiment with different settings if any modifications to the current system are introduced.
  10. Uncheck “Auto Calibrate” if data are only collected over 24-hr period or less (**Figure 6H-2**).

### 5. Daily procedures for lickometer-equipped two-bottle choice home cage drinking study

NOTE: Follow the steps outlined in **Section 3** for standard 2BC with the following modifications.

1. On Mondays, Wednesdays, and Fridays, before putting on the MWF bottles, use the sipper blockers to cover the lickometer sipper openings on all cages (see **Section 4: step 5.7** and **Figure 5E**).
2. Before recording, select the correct “Experimental Side” on the main GUI page, which will be the side of the EtOH bottle for that day (**Figure 6G**).
3. After installing all MWF bottles, tap “START!” on the main page (**Figure 6G**) to start calibrating sensors, during which users are recommended to take a step away from the system and make sure rats are not licking the sippers (**Figure 6I-1**). Calibration takes a few seconds to complete, after which the system enters the main recording page showing the cumulative lick numbers on each bottle for each lickometer (**Figure 6I-2**).
4. Raise all sipper blockers above the cage top to allow access to the sippers (see **Section 4: step 5.8** and **Figure 5F**), and gently return all filter tops as described (see **Section 4: step 4.6** and **Figure 4F**).
5. Tap “Refresh” to update the lick numbers on screen, which will not update in real time by itself (**Figure 6I-2**).
6. On Tuesdays, Thursdays, and Saturdays, tap “Save & Quit” to stop recording and return to the main page (**Figure 6I-2**), before removing the MWF bottles
7. To remove SD card for data transferring, tap “Eject SD” before unplugging the SD card (**Figure 6G**). Insert the SD card into any computer and copy/paste the .csv data file named after the bottle ON date to that computer. To remount the SD card for the next recording, insert the card into the interface and tap “Mount SD” (**Figure 6J**). NOTE: We recommend backing up data files daily.
8. Limit body weight measurements to once a week and perform this task on the day designated for cage change in order to minimize unnecessary disturbance to the lickometers and sensors. Unplug all male lickometer cables from the matching female sensor cables before moving any cages. After weighing each rat, return the rat to a clean cage. Transfer the lickometer to a new cage top and reconnect each lickometer to the corresponding sensor when necessary.

### 6. Data analysis

1. Prior to evaluating dependent measures, researchers should correct intake data for spillage that inevitably occurs as a result of changing bottles each day. To do so, the change in bottle weight for EtOH and H_2_O bottles on the spillage cage should be averaged over the course of the entire study period. Any values greater than two standard deviations from the mean for each bottle should be excluded. After excluding these outliers, the calculated means should be subtracted from the changes in EtOH and H_2_O bottle weight obtained from bottles on cages containing animals. By subtracting this amount, researchers can be confident that average loss resulting from spillage of a given solution is not considered in measures of consumption. Note that subtraction of the average spillage amount may result in a negative value for some animals on some occasions. These values should be converted to zero and interpreted as absence of any intake for that solution.
2. The primary dependent measures for two-bottle choice drinking experiments are EtOH intake (g/kg) and preference (%). Researchers can calculate intake from change in EtOH bottle weight (g) and normalize to body weight (kg) using the following steps:
  1. Convert change in bottle weight (g) to change in volume (mL) using the following equation:

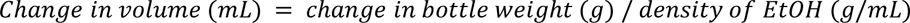 NOTE: The density of pure ethanol is 0.789 g/mL. Therefore, the contributing density of 20% EtOH is 0.1578 g/mL. H2O weighs 1 g/mL. Therefore, the contributing density of 80% H2O = 0.80 g/mL. Consequently, every mL of a 20% EtOH solution (v/v) is comprised of 0.1578 (EtOH) + 0.80 (H2O) = 0.9578 g/mL. Therefore, the final equation when using 20% (v/v) EtOH concentration is:

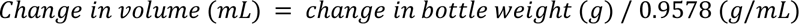
  2. Calculate the dose of ethanol consumed using the equation below. Use body weights obtained during the same week as the intake data in order to calculate ethanol consumed in terms of animal body weight.

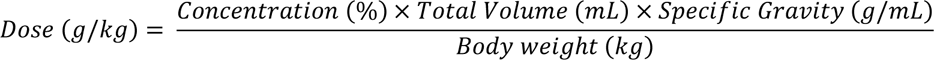
  3. Calculate preference using the following equation:

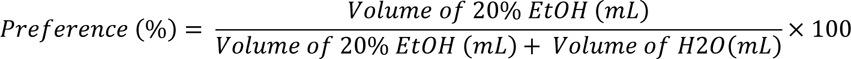
3. Like its mouse counterpart^27^, LIQ HDR captures a variety of drinking microstructure variables including 1) lick duration, 2) bout number, 3) bout duration, 4) bout lick number, and 5) bout lick duration. These measures can subsequently be used to calculate the 6) average individual lick duration, 7) average individual bout duration, 8) average number of licks per bout, 9) average lick frequency, 10) estimated inter-lick interval, and 11) estimated inter-bout interval. The Arduino interface provides these data at 1-min resolution in a .csv file. Researchers can use the analysis app provided with the original LIQ HD protocol^27^ or their preferred post-processing platform to collate and summarize data according to research needs.

## REPRESENTATIVE RESULTS

### 1. Standard intermittent two-bottle choice home cage EtOH drinking data

Using the standard procedure (**Figure 7A**), researchers can capture 24 hr intake (**Figures 7B-D**) as well as binge-like drinking if bottle weights are collected shortly after the onset of the drinking session (**Figures 7E-F**). By calculating ethanol intake relative to water intake researchers can also obtain preference data at all time points measured (**Figures 7D&F**). As shown in **Figure 7C**, rats frequently exhibit a period of escalation in ethanol intake over the course of the first few days of two-bottle choice access. This is followed by a plateau during which intake remains relatively stable. Although consistently observed in male rats^10, 29–33^, this pattern is not always observed in females. Instead, some studies report stable levels of ethanol intake throughout the study period in female rats^34, 37^. In addition, some studies suggest that ethanol intake may escalate further with more prolonged periods of access^33, 35^. Daily intake can vary significantly across individuals. However, average daily has been well-characterized for the strains most commonly used in alcohol research and can be used as a benchmark for success when looking at grouped data^10, 11, 30–32, 35^.

**Figure 7:**
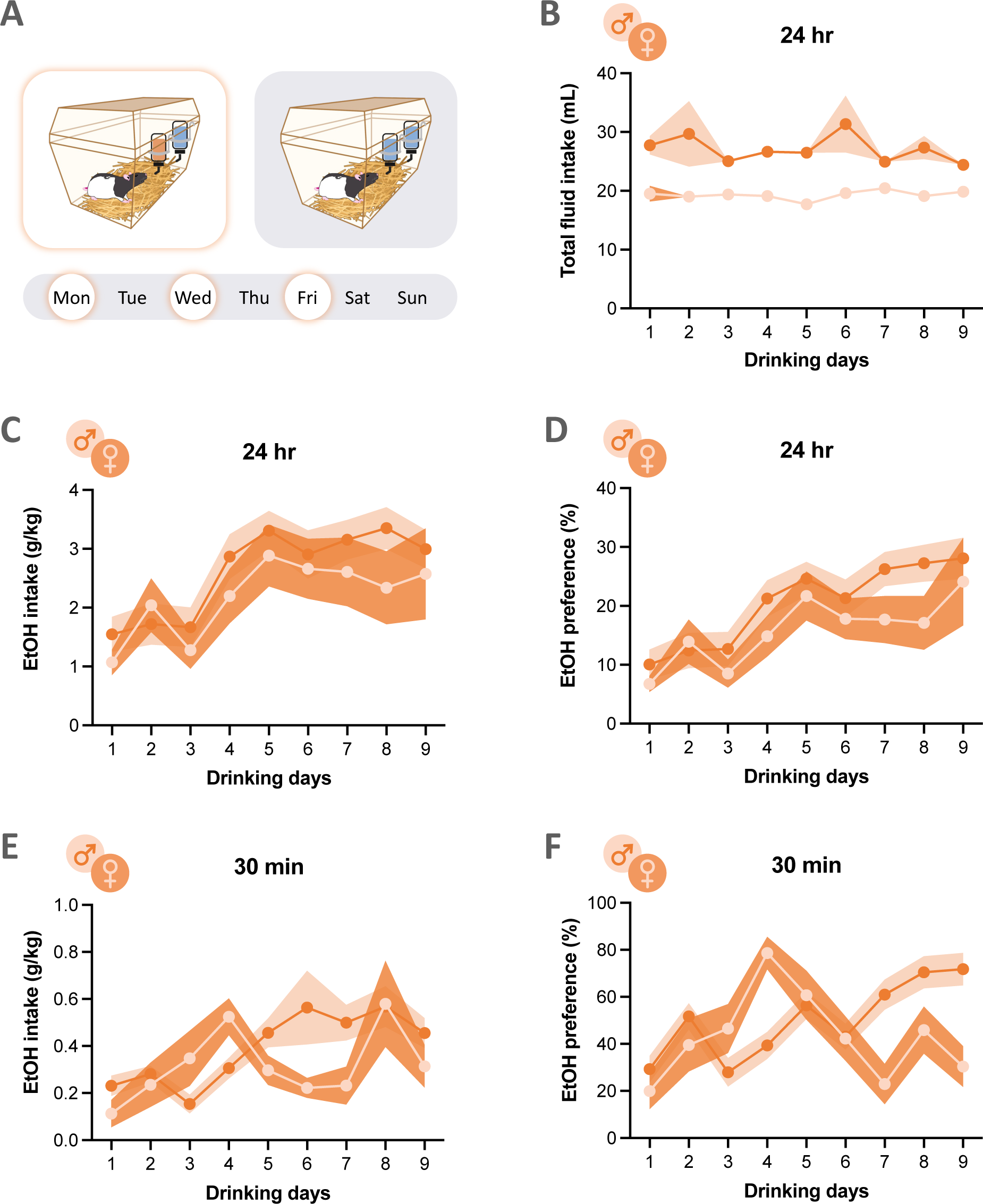
Representative two-bottle choice drinking data captured using the standard procedure. In the intermittent-access two-bottle choice home cage EtOH drinking paradigm **(A)**, rats receive 20% (v/v) EtOH solution (orange) and H_2_O (blue) on Mondays, Wednesdays, and Fridays (MWF), and two bottles containing H_2_O on Tuesdays, Thursdays, and weekends. Measures of daily fluid intake reveal that while rats maintain a relatively consistent level of total fluid intake (mL) throughout the study period **(B)**, rats often exhibit escalation in EtOH intake over the course of the first few drinking sessions after which intake levels plateau **(C)**. This pattern of escalation and maintenance is similarly reflected in preference for EtOH **(D)**. Researchers can also capture binge-like EtOH intake **(E)** if bottle weights are collected after the first 30-60 min of session start. This is often associated with high preference for EtOH over H_2_O **(F)**. Shaded error bars represent ±SEM.

As show in **Figure 7E**, many rats consume a significant portion of their daily ethanol intake shortly after the onset of the drinking session^11, 30^. This binge-like pattern of ethanol intake is often associated with significant preference for ethanol over water (**Figure 7F**). Researchers should not be surprised, however, if 24-hr preference does not exceed 50%. This is particularly true for outbred or alcohol non-preferring rodent strains and studies limited to relatively short periods of two-bottle choice access.

### 2. Validation of LIQ HDR for two-bottle choice drinking microstructure analysis

Addition of the LIQ HDR system to the standard rat two-bottle choice paradigm allows researchers to accurately and precisely predict intake using captured drinking data. This is confirmed by the presence of a strong and significant correlation between the cumulative number of licks detected and change in bottle weights for both the EtOH (R^2^ = 0.7789, F_(1,113)_ = 398.1, p < 0.0001) and H_2_O bottle (R^2^ = 0.7910, F_(1,117)_ = 442.9, p < 0.0001) over a 24-hr period (**Figure 8A**). This is also true for days when rats received two bottles containing H_2_O (R^2^ = 0.8875, F_(1,188)_ = 1484, p < 0.0001) (**Figure 8B**).

**Figure 8:**
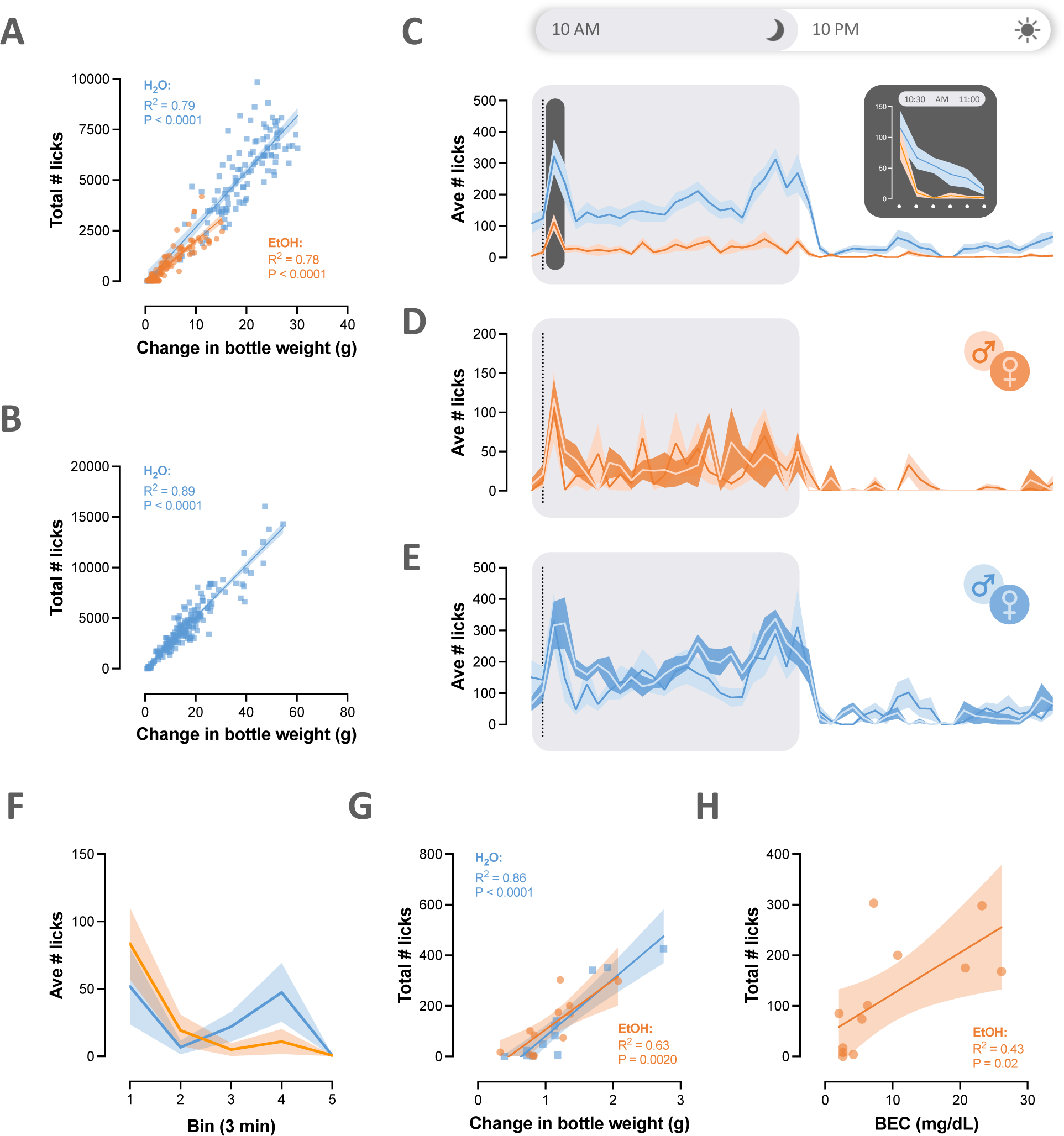
LIQ HDR accurately captures consumption at high temporal resolution. The precision and accuracy of the LIQ HDR system is confirmed, in part, by the presence of a significant positive correlation between number of licks and change in bottle weight on days when rats receive H_2_O and EtOH **(A)** as well as days when both bottles contain H_2_O **(B)**. Differences in drinking patterns between H_2_O (blue) and EtOH (orange) across the 24-hr drinking period can be easily assessed by binning average licks into 30-min intervals **(C)**. Inset corresponds to average licks in 5-min bins during the first 30 min of bottle access (highlighted in gray) during which time rats often exhibit significant front-loading behavior. Between-group differences in drinking patterns can also be examined using a similar approach as exemplified by lick microstructure for EtOH **(D)** and H_2_O **(E)** depicted for male and female rats across a 24-hr period. Drinking patterns can also be assessed in high temporal resolution across shorter testing period **(F)** during which LIQ HDR continues to accurately track intake **(G)** and predict blood ethanol concentration (BEC) even for relatively low levels of EtOH intake **(H)**. Shaded error bars represent ±SEM.

LIQ HDR also enables precise analysis of drinking patterns at high temporal resolution that is not afforded by the standard two-bottle choice paradigm. Using this approach, researchers have the opportunity to examine differences in drinking patterns between different solutions, across the light cycle, between sexes, drug groups, and more. To illustrate, binning average licks into 30-min intervals allows for easy analysis of average 24 hr drinking patterns. As expected, this analysis reveals significantly higher intakes for both EtOH and H_2_O during the dark cycle, when rats are awake, than the light cycle, when rats are typically sleeping (**Figure 8C**). Using a similar approach, researchers can explore group differences in drinking patterns. For example, drinking microstructure can be compared between male and female rats by examining the average number of licks for EtOH (**Figure 8D**) or water (**Figure 8E**) across a 24 hr period. To investigate front-loading and/or binge-like drinking, researchers can increase the resolution of their analysis to focus on drinking occurring shortly after session onset using a smaller binned window to summarize data. Doing so allows researchers to investigate periods of binge-like drinking without requiring collection of multiple bottle weights within a 24-hr period. As shown in **Figure 8C** (inset), this analysis reveals that most of the drinking captured by a bottle weight obtained 30 min after session onset occurs in the first five minutes, rather than throughout the 30-min period. Data such as these can aid in the identification of optimal time points for the collection of blood samples to measure blood ethanol concentration (BEC). Indeed, a similar pattern of front-loading was observed on a separate drinking day during which high-resolution drinking patterns were captured in a period as short as 15 min (**Figure 8F**). Average licks recorded during this session reliably predicted change in bottle weight (EtOH: R^2^ = 0.6316, F_(1,10)_ = 17.15, p = 0.0020; H_2_O: R^2^ = 0.8601, F_(1,10)_ = 61.47, p < 0.0001; **Figure 8G**). Moreover, BECs taken from blood samples obtained 15 min after session onset correlated well with change in bottle weight (R^2^ = 0.4327, F(1,10) = 7.627, p = 0.0201; **Figure 8H**) even for relatively low levels of EtOH intake.

LIQ HDR captures various measures of drinking microstructure which are detected and calculated in real-time during drinking sessions and saved in 1-min bins similar to lick detection. These measures are briefly outlined in Protocol **Section 6: step 3** and defined previously^27^. Researchers can conduct detailed analyses of changes in individual drinking microstructure and examine how and to what degree drinking patterns contribute to differences in overall intake. To illustrate, we found that although male and female rats exhibited a similar number of licks, drinking bouts, and bout length (**Figure 9A-C**) when drinking water, this was associated with greater overall intake (g/kg) in females than males (t-test; t=2.882, df=10, p=0.0163; **Figure 9D**). This is expected given the sexual dimorphism in body weight which presumably facilitates greater levels of intake by body weight for females than males for an equivalent number of licks. Interestingly, this is in contrast to comparison of sex differences in microstructure patterns for EtOH, which revealed similar drinking microstructure associated with similar levels of EtOH intake (**Figures 9E-H**). These data suggest that while lick volume for water is similar between sexes, compared to males, female rats have smaller lick volume for EtOH. Together, these lickometer microstructural data support the idea that male and female rats likely differ in voluntary drinking strategies based on the type of fluid consumed in the home-cage setting.

**Figure 9:**
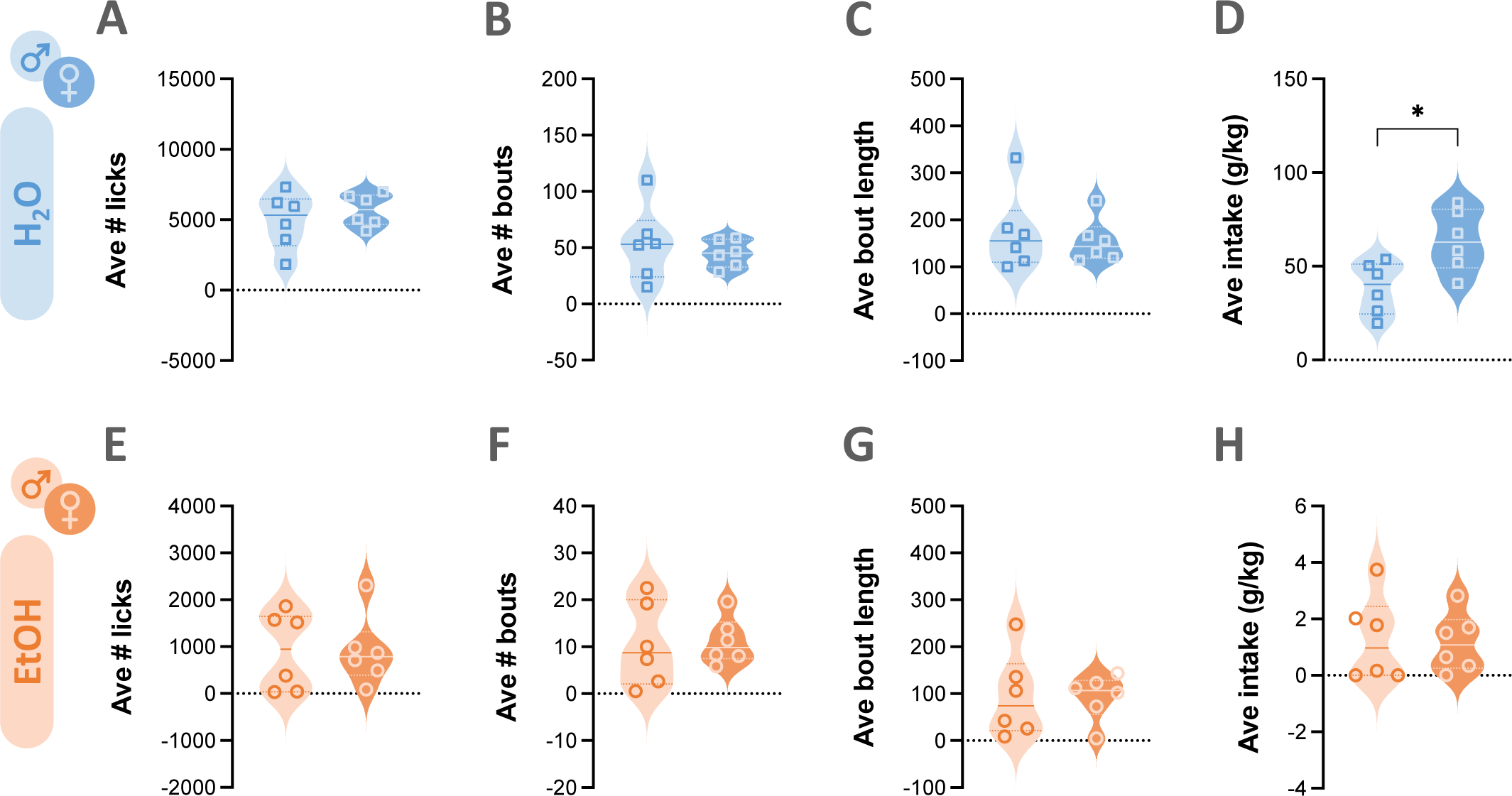
Representative drinking microstructure data. LIQ HDR enables capture of measures of drinking microstructure including the number of licks **(A)**, number of drinking bouts **(B)**, and bout length **(C)**, in addition to measuring total water intake by change in bottle weight **(D)**. Similar measures can be obtained for ethanol **(E-H)**. Between-group comparisons of these measures can identify distinct drinking patterns as is the case in the current example, which reveals sex differences in drinking strategies based on the solution consumed. Solid line indicates median and dotted lines indicate quartiles. \**p* < 0.05.

## DISCUSSION

The current protocol provides step-by-step instructions for capturing ethanol drinking data using an intermittent-access two-bottle choice home cage procedure. This method can be implemented with relative ease and with little-to-no cost or need for specialized research equipment. Using the procedures provided here for constructing and implementing LIQ HDR – a lickometer system for use in rats – researchers can capture high-resolution drinking microstructure data in addition to standard measures of intake and preference.

The intermittent-access schedule described in the current protocol is a widely used procedure in large part because it is well-known to engender higher levels of ethanol intake than many alternative schedules of ethanol access. Indeed, previous research has shown that rats with access on an intermittent schedule have higher average daily ethanol intake than rats with continuous 24h access^9, 10^. Nevertheless, different schedules of ethanol access may be desirable depending on the research question or other experimental design parameters. Researchers can very easily modify the current protocol to accommodate different schedules (e.g., continuous access, drinking in the dark, etc) by simply changing the bottle on/off time and days.

Although two-bottle choice drinking is a relatively simple to implement paradigm, issues can arise that have the potential to compromise data collection. It is, therefore, crucial that experimenters inspect data on a daily basis in order to quickly identify potential problems. In particular, researchers should pay close attention to very large or very small changes in bottle weights that may be indicative of a leaky or clogged sipper, respectively. Catching signs of these issues early can prevent exclusion of all data from a given subject from analysis. Indeed, if researchers notice drastic changes (or absence of change) in bottle weight over a 24h period, a simple switch to a new bottle can help to confirm whether a given bottle should be removed from circulation. For these reasons, it is wise to have extra bottles and sippers on hand. However, it should be noted that some rats naturally exhibit very low levels of ethanol drinking^e.g., 32^. This is particularly true for rats of certain strains^see 11 for review^. Thus, researchers should not be alarmed if a small population of animals in a given experiment have low or no ethanol consumption. Likewise, some animals are prone to “play” with the sipper/bottle in ways that can promote spillage. Although rare, repeated instances of high changes in bottle weight that are not associated with consumption may require that a particular animal be removed from the experiment.

Our detailed step-by-step protocol demonstrates the ease of implementation and use of the LIQ HDR lickometer system for rats. Researchers with no prior knowledge or skills in microcontroller electronics and programming can readily adapt this system to their experimental needs with little or no modification. Our representative data further demonstrates its successful application for rat home cage drinking studies and its powerful capacity to capture numerous drinking microstructure measures at high temporal resolution. LIQ HDR includes several modifications to the lickometer design relative to its mouse predecessor^27^. This was done not only as a necessity for successful adaptation for use in rats, but also to improve functionality and user experience. In-cage components including the 3D printed bottle holder, sipper blocker, and electronic components are relocated to the cage top to protect against chewing, which can result in frequent and significant data loss and hardware turnover. The addition of a laser-cut acrylic panel further helps prevent destruction of these vulnerable components. In contrast to the relatively labor-intensive custom-made bottles used in the LIQ HD design, LIQ HDR’s 3D printed bottle holder is designed to be compatible with commercial off-the-shelf rodent drinking bottles. Bottles are installed on the cage top and accessible to rats in much of the same way as drinking bottles are applied during standard care and husbandry. The addition of sipper blockers is a simple but effective solution that improves system reliability and prevents valuable data loss that, in the absence of the blocker, can result from drinking that occurs prior to initiating data capture on the Arduino interface or erroneous lick measures that occur when rats interact with the sippers during daily calibration. While the considerably larger rat cages necessitate scaling up the original hardware design in size, the use of significantly longer cables can negatively impact the sensitivity and accuracy of capacitive sensing due to interference, as well as the integrity of I^2^C communication between the daisy-chained sensor boards and Arduino interface. We circumvent this issue with the adoption of the QwiicBus communication system to achieve ultra long-range, near lossless signal transmission via Ethernet cables.

The reliability and effectiveness of any lickometer system requires proper construction and installation as well as extensive testing, diligent error monitoring, and effective troubleshooting. All DIY components need to be made and assembled with skill and care. This is especially important for steps involving soldering, which may require practice for beginners. Regardless, extensive testing of all lickometers is required before use in actual experiments. If the system must be modified in order to adapt to individual experimental needs, the sensor settings (e.g. touch and release thresholds) may need to be further fine-tuned for accurate lick detection. For first-time users, we suggest testing the system in a brief, small-sample, pilot study using a highly palatable solution that will engender high levels of drinking. Lick detection can be verified in real-time by visually inspecting the sensor board as the LED flashes whenever a lick is registered. Strong, significant, and consistent correlations between lick number and change in bottle weight should be used as a benchmark for proper hardware and software setup. Even after the setup is finalized, we recommend initiating recording for a given experiment on a day when rats receive two bottles containing water in order to troubleshoot equipment issues that may be evident at the beginning of an experiment. It may also be advantageous to test lickometer function on the water day associated with cage change since this requires that lickometers be disconnected and reconnected to the sensor. Random failure of individual lickometer or sensor, which would result in very obvious outlier data (e.g. no or few licks detected despite substantial change in bottle weight) consistently for the same lickometer, is indicative of faulty component(s). We recommend preparing extra components for each element of the lickometer system in advance to enable easy replacement as necessary. Finally, the importance of making sure that sensor cables do not overlap or bend excessively cannot be overemphasized. Doing so can result in random instances of poor lick detection evident by an unreasonably low lick number for a given change in bottle weight or extremely large lick durations (close to 60,000 ms for a 1-min bin) spanning minutes or hours. Nevertheless, small losses of data are inevitable in studies that collect data 24h/day for many animals over many days using either the standard or lickometer-equipped approach. Researchers can exclude erroneous data for the rare drinking sessions when bottle weights and/or licks are clearly not reflective of intake (e.g., sipper clog, calibration error, etc). On occasions such as these, the average value for a given measure (e.g., intake (g/kg), licks, etc) across the study period for that particular subject can be substituted for the missing value during data analysis.

One of the advantages of the two-bottle choice drinking paradigm is the voluntary aspect of ethanol intake. This affords excellent face validity particularly in light of findings showing that experimenter administered alcohol can produce significantly different effects on brain function and behavior compared to voluntarily self-administered alcohol^36^. Moreover, the voluntary nature of the paradigm allows researchers to capture of individual differences in intake similar to the variability in alcohol preference and drinking patterns observed in humans. These individual differences can provide a powerful opportunity for researchers to correlate consumption (intake and/or patterns) with the magnitude of pathological effects observed.

Nevertheless, some limitations of the two-bottle home cage drinking paradigm should be considered. For example, although individual variability in ethanol intake can be advantageous for some experiments, it may be a disadvantage in other experiments where a consistent level of intoxication or ethanol exposure may be desirable. This is particularly true for experiments examining the consequences of dependence and withdrawal from chronic ethanol exposure. Importantly, two-bottle choice drinking is not characterized as a model of dependence – even in rats with access to ethanol for prolonged periods of time. Thus, in the absence of accompanying symptoms of withdrawal, researchers should use caution in interpreting the consequences of prolonged ethanol consumption as effects of dependence and/or withdrawal. Indeed, it is possible, if not plausible, that rats drinking relatively large quantities of ethanol during the two-bottle choice procedure exhibit symptoms of withdrawal but that rats drinking lower total volumes in a given 24 hr period do not show similar signs of dependence. Drinking patterns can also play an important role in this consideration as two animals drinking the same amount over a 24 hr period can experience vastly different levels of intoxication (i.e., BEC) depending on the pattern of their ethanol intake^7^. This highlights the utility of LIQ HDR in the two-bottle choice procedure as it can readily identify rats that consume a given dose of ethanol at a fast rate (i.e., gulp) versus those who consume the same dose at a much slower rate (i.e., sip). Although potentially undesirable for studies investigating effects of a controlled dose of ethanol, detection of changes in drinking patterns (e.g., reduced binge episodes) following a particular experimental intervention have significant translational value given the harm reduction approach that is increasingly pursued in the clinic. Finally, effort is not required in order for animals to gain access to ethanol in home cage drinking paradigms including the one described here. Therefore, the utility of this paradigm is limited to analysis of drinking alone whereas operant methods are favored for instances when researchers are interested in examining alcohol seeking – an important feature of AUD.

In summary, the protocol described here provides researchers with all the information and resources necessary to measure ethanol intake in rats in the home cage setting. We present options for relatively low- and high-resolution approaches providing researchers with the flexibility to choose the method that is most suitable to their research needs.

## ACKNOWLEDGEMENTS

The authors thank Joseph Pitock and Katie Przybysz, PhD for technical assistance during LIQ HDR development. We also thank Nicholas Petersen and Marie Doyle, PhD for many helpful conversations during the development of LIQ HDR. This work was supported by the National Institute on Alcohol Abuse and Alcoholism at the National Institutes of Health (P50 AA022538 and R01 AA029130 to EJG).

---

**Table of Materials.**
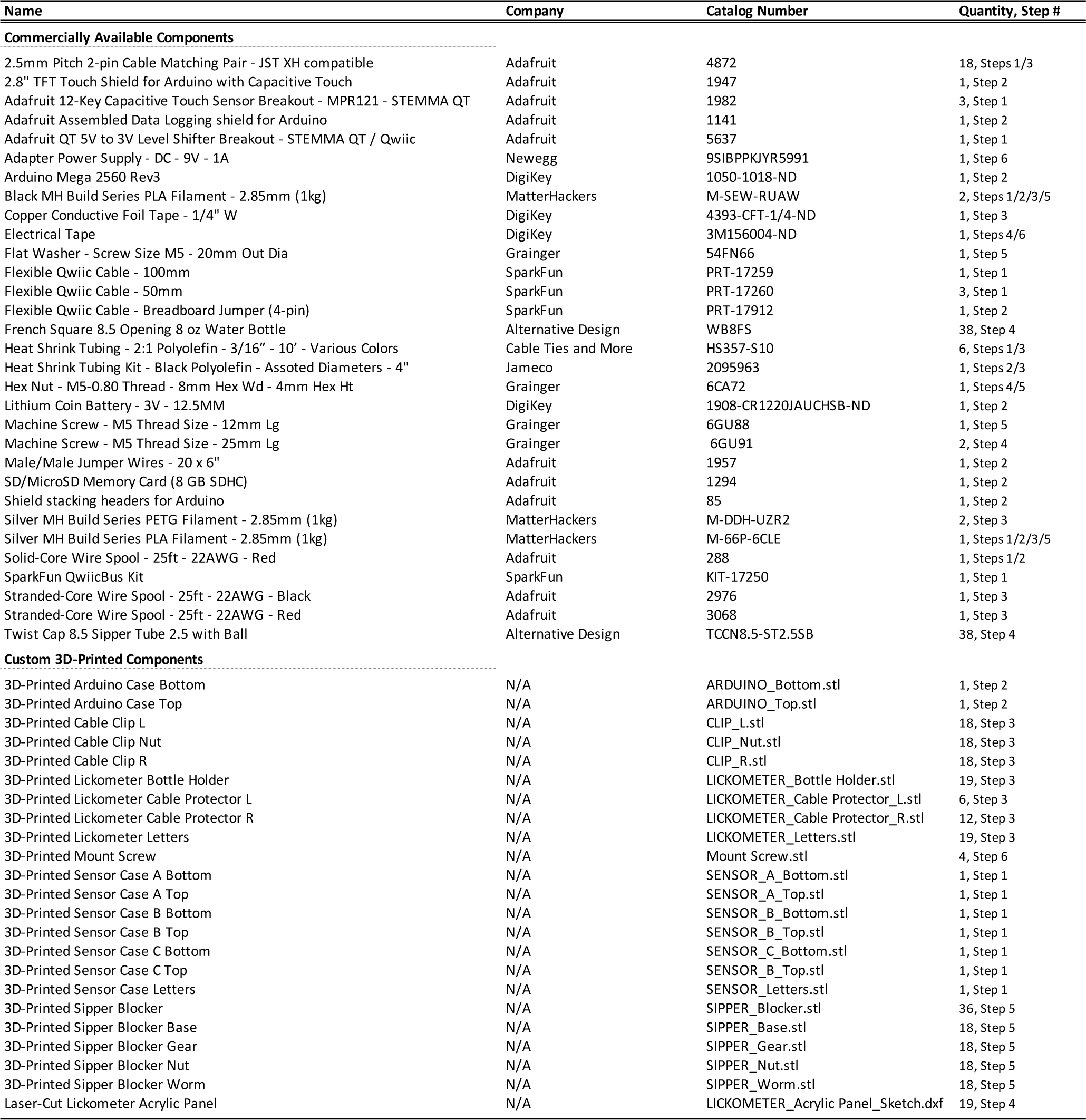
All materials required to equip n=18 cages with LIQ HDR for two-bottle choice experiments.

